# Integrin-linked kinase is a key signal factor involved in Nogo-66-induced inhibition of neurite outgrowth

**DOI:** 10.1101/2020.04.14.040444

**Authors:** Ya-ping Yu, Qiang-ping Wang, Jian-Ying Shen, Nan-xiang Xiong, Hua Yu, Peng Fu, Lei Wang, Ye Yuan, Hong-yang Zhao, Fang-cheng Zhang, Hendrik Pool

**Affiliations:** Department of Neurosurgery, Union HospitalTongji Medical College, Huazhong University of Science and Technology, Wuhan 430022, China; Department of Anatomy, Section of Histology and Embryology, Tongji Medical College, Huazhong University of Science and Technology,Wuhan,430030, China; Department of International Education, Tongji Medical College, Huazhong University of science and Technology, Wuhan 430030, China

**Author notes:** Ya-ping Yu, Qiang-ping Wang and Jian-Ying Shen contributed equally to this work. Correspondence to Prof. Nan-xiang Xiong. Address: Dept. of Neurosurgery, Union Hospital, Tongji Medical College, Huazhong University of Science and Technology, Wuhan, China, 430022.

**Keywords:** Integrin-linked kinase, Nogo-66, Protein kinase B, Glycogen synthase kinase-3β, tau, neurite outgrowth

## Abstract

Nogo-66, the extracellular domain of Nogo-A, has been identified as the most important myelin-associated neuronal growth inhibitor. Evidence suggested that Nogo-66 exert its neurite inhibition effect via a Nogo-66/Protein kinase B (PKB)/Glycogen synthase kinase-3β (GSK-3β)/tau signaling pathway. Integrin-linked kinase (ILK) is a serine/threonine kinase mediating axon upstream growth of PKB and GSK-3β. However, the contribution of ILK to the Nogo-66-induced inhibition of neurite, is not clear. In this study, we set out to reveal the role of ILK on Nogo-66 signaling *in vitro* and *in vivo*. To deteremine this directly, Recombinant adenoviruses were constructed to upregulate or downregulate the expresioon of ILK in Neuro 2a (N2a)and analysis the change of downstream molecule and neurite length. The results showed that Nogo-66 inhibited the phosphorylation of ILK, while ILK regulated the phosphorylation of PKB and GSK-3β, and the expression of tau in Nogo-66-treated N2a cells. ILK overexpression through lentivirus vector transfection reduced the inhibitory effect of neurite outgrowth induced by Nogo-66 in cortical neurons. The Tau expression in the complete spinal cord transection rat model was promoted by the overexpression of ILK. Our findings indicated that ILK is a key signal factor involved in Nogo-66-induced inhibition of neurite outgrowth. The mechanism of Nogo-66 signaling pathway was further explained and a proper target for the promotion of neural regeneration was also provided by this study.

## Introduction

The growth-inhibitory environment induced by myelin inhibitors has been identified as a key factor contributing to the failure of injured axons to regenerate in the adult mammalian central nervous system (CNS)^[1, 2]^. The main validated myelin inhibitors included neurite outgrowth inhibitor (Nogo-A), oligodendrocyte-myelin glycoprotein (OMgp), myelin-associated glycoprotein (MAG) and chondroitin sulfate proteoglycan (CSPG)^[3, 4]^. Among these, Nogo-A has been identified as the most important myelin inhibitor^[5]^. Nogo-66, the inhibitory domains of Nogo-A, is a 66-residue extracellular domain of Nogo-A, which exerts growth inhibition via binding to multiple neuronal receptors, including the paired immunoglobulin-like receptor B, the ligand-binding Nogo-66 receptor 1 (NgR1) and other co-receptors, and then transducing the inhibitory signal to the cell interior^[5]^.

Protein kinase B (PKB) is a protein serine/threonine kinase that plays a dominant role in cellular homeostasis and intracellular signaling^[6]^, while Glycogen synthase kinase-3β (GSK-3β), which is the most important downstream molecule of PKB, implicated in polarity, specification, differentiation and axon growth in the nervous system through phosphorylating several microtubule associated proteins^[7]^. Previous studies found that Nogo-66 regulated the activity of PKB and GSK-3β through reducing the phosphorylation at Ser473 and Ser9 respectively^[8, 9]^. Tau, a microtubule-associated protein, plays a critical role in microtubule assembly and stabilization^[10]^. Accumulating evidence has indicated that tau is a key factor closely related to axonal elongation and neurite outgrowth, and that it carries out its physiological functions on the status of phosphorylation^[11]^. Research found that the phosphorylation of tau was regulated by GSK-3β^[12]^. This evidence indicated that Nogo-66 exerted its neurite inhibition effect via a Nogo-66/PKB/GSK-3β/tau signaling pathway way. However, the underlying cytokines and upstream mechanisms of PKB/GSK-3β have not yet been fully defined.

Integrin-linked kinase (ILK) is a serine/threonine kinase which has recently been implemented in axonal growth^[13]^. Its downstream function of growth factor receptors and β-integrins to regulate cell differentiation and migration^[14]^. Additionally, PKB and GSK-3β are the main targets currently known to mediate the effects of ILK-mediated axon downstream growth of ILK. However, whether ILK is involved in Nogo-66-induced inhibition of the neurite outgrowth signaling pathway, is still unknown. In the present study, the role of ILK in Nogo-66-induced inhibition of neurite outgrowth *in vitro* and *in vivo* were studied.

## Experimental procedures

The ethics committee of Tongji Medical College, Huazhong University of Science and Technology granted the approval to conduct this study. All experiments were carried out in accordance with the guidelines for molecular and animal research at Tongji Medical College.

### Antibodies and Reagents

The main primary antibodies and compounds: Nogo-66, 0.5 mg (Bioss, Beijing, China). Anti-integrin-linked ILK antibody, ab52480, rabbit (1:1000; Abcam, Cambridge, UK); Anti-Integrin linked ILK (phospho S246), ab111435, rabbit (1:1000; Abcam, Cambridge, UK); Anti-GSK3β antibody, ab32391, rabbit (1:1000; Abcam, Cambridge, UK); Anti-GSK3β (phospho S9) antibody, ab75814, rabbit (1:1000; Abcam, Cambridge, UK); Anti-AKT1 (phospho S473) antibody, ab81283, (1:1000; Abcam, Cambridge, UK); Anti-tau (phospho S262), mouse (1:200; Abcam, Cambridge, UK); glyceraldehyde-3-phosphate dehydrogenase (GADPH), mouse (1:200; Abcam, Cambridge, UK); dimetilsulfóxido (DMSO; Amresco, Solon, USA). The main secondary antibodies: Goat Anti-Rabbit IgG (Alexa Fluor 647), ab150079, (1:800; Abcam, Cambridge, UK); Goat Anti-Rabbit IgG (FITC) (ab6717); rabbit anti-goat IgG BA1060 (1:200; Boster, Wuhan, China); rabbit anti-goat IgG-Cy3 (1:50; Boster, Wuhan, China); goat anti-mouse IgG-Cy3 (1:50; Boster, Wuhan, China);

### Cell cultivation

Within 24 hours of birth, primary cerebral cortical neurons were isolated from newborn Sprague-Dawley (SD) rats (Experimental Animal Center of Third Military Medical University, Chongqing, China). The procedures for isolation and cultivation were the same as previously described^[12]^. Due to its neuron-like property, the Neuro 2a (N2a) cell line (a gift from Prof. Honglian Li) was selected for this research. The cells were cultivated on collagen-coated dishes in Dulbecco’s modified Eagle’s medium (DMEM)/F12 (Gibco, New York, USA), supplemented with penicillin-streptomycin solution (Biyuntian Biotechnology, Shanghai, China), 5% fetal bovine serum (FBS; Zhongshan Biotechnology, Beijing, uChina), and 2 mM l-glutamine (Gibco, New York, USA). Cells were treated with Nogo-66 (15 ng/mL, Biosynthesis Biotechnology, Beijing, China) for a staggered time schedule (0min, 5min, 30min, 2h, 6h, 24h and 48h), followed by cultivation with normal medium.

### Recombinant adenovirus construction and infection

The overexpression (Ove.) and interference (Int.) ILK plasmids were purchased from Shandong Vigene Bioscience, Jinan, China. ILK sequences were selected and constructed into a lentivirus vector (PLent-GFP-Puro-CMV) for the overexpression of ILK. The ILK knockdown was performed with the following siRNA:5□-GACAACACAGAGAACGACCTCAATTCAAGAGATTGAGGTCGTTCTCTGTGTTGTCTTTTTT-3□. This sequence was constructed into a pLent-CMV-GFP-Puro vector to construct a lentivirus vector expressing small hairpin RNA (shRNA) targeting ILK. Cells transfected with blank vectors (Bla.) were used as controls. These constructs were verified by polymerase chain reaction (PCR) and western blot analysis. The resultant recombinant adenoviruses were plaque-purified three times, expanded, and tittered. For cell infection, recombinant adenoviruse were added to the culture medium, and 15-20 hr later, approximately half of the medium was replaced with fresh medium including nogo-66 (15 ng/mL), and those target cells were harvested after staggered time schedule described as above.

### Protein isolation and western blot analysis

Cells were rinsed four times with ice-cold PBS before addition of SDS lysis buffer.Western blot analysis was used to detect the expression level of ILK, phosphorylated ILK at residues Ser246 (p-ILK), PKB, phosphorylated PKB at residues Ser473 (p-PKB), GSK-3β, phosphorylated GSK-3β at residues Ser9 (p-GSK-3β) and tau. Cell or tissue samples were homogenized in a lysis buffer and electrophoresed in a 10% sodium dodecyl sulfate-polyacrylamide gel electrophoresis (SDS-PAGE) gel at 100 V for 2 h. Fractionated proteins were then electroblotted onto a polyvinylidene difluoride (PVDF) membrane (Bio-Rad, Hercules, USA) and incubated with specific antibodies as internal control. Image J were used to detect the band densities (Bio-Rad, Hercules, CA).

### Immunofluorescence staining and measurement analysis

N2a cells were washed with ice-cold polybutylene succinate (PBS) and then fixed with 4% paraformaldehyde for 15 min. Subsequently, cells were permeabilized with 0.1% Triton X-100 for 30 min and incubated with primary antibodies which were diluted in 10% normal goat serum and 1% Triton X-100 in PBS for 10 min. Specimens were then incubated with secondary antibodies for 1h at room temperature, followed by three times washes using PBS. A laser scanning confocal microscope (Olympus Fluoview FV1000) was used to capture images.Then Image J were used for the measurement of immunofluorescence intensity.

#### Quantitative analysis of neurite outgrowth

A digital phase-contrast inverted microscope (Olympus, Tokyo, Japan), with a charge-coupled device camera was used to analyze the morphological changes of axons and the Image J software was used to measure the neurite length. The neurite outgrowth from N2a were quantitatively measured after optimization of culture conditions. After cultures were fixed, microscope images were captured and an image analysis algorithm was developed using Image-Pro Plus software to allow quantitative analysis. Extenision longer than 15μm (longer that one cell diameter) were considered a neurite. For every group, both transfected cells and surrounding nontransfected cells were measured and compared. Average neurite length of each group were calculated. Measurements were made on duplicated wells (n= 6), and each experiment was repeated at least twice.

### Complete spinal cord transection model of rats

Adult female SD rats (Experimental Animal Center of Third Military Medical University, Chongqing, China), weighing between 250~300g were used to construct the model. Rats were anesthetized using pentobarbital with a dose of 40 to 50 mg/kg.. A laminectomy was performed once the 9^th^ and 10^th^ thoracic vertebrae were exposed. A micro scissor was then used to completely transect the spinal cord. The muscles and skin were sutured, concluding the surgery. The sham-operated group only received muscle and skin incisions and sutures, whereas the control group didn’t undergo any surgical procedures. These models were executed at a set time point and the injured spinal cord tissues were obtained for experiments.

The interventional experiment was performed seven days after the injury. After anesthetization, the rats were injected with 1μL of saline (control group), EGFP-Lenti (Bla. group), ILK-Lenti-Ove (Ove. group) and shRNA ILK-Lenti-Int (Int. group) at the site of the sensorimotor cortex (coordinate from bregma in cm: 0.5/1.5; depth: 1.5 mm). A microinjection pump fixed on a stereotactic instrument (Stoelting, USA) was applied using with this procedure. The detailed procedures were the same as were described previously^[12]^. Seven days after the transfection, injured spinal cord tissue were harvested. Three rats in each group were used for this experiment.

### Statistical analysis

The Statistical Package for the Social Sciences version 19.0 software (SPSS, Chicago, IL, USA) was used for statistical analysis. Data is expressed as mean ± standard deviation (SD). The *t-*test was applied to compare the differences among groups. The time effects of Nogo-66 and ILK were detected using One-way analysis of variance (ANOVA) with Tukey’s post-hoc analysis. P < 0.05 was considered to be statistically significant.

## Results

### Nogo-66 Downregulated Phosphorylation Level of ILK in N2a Cells

Determining whether Nogo-66 regulates ILK signaling in N2a cells, the expression levels of p-ILK and total ILK were detected after exposure to Nogo-66 treatment for a variety of time durations. Our results showed that Nogo-66 attenuated p-ILK in a time-dependent manner (Fig. 1A), and statistical differences were observed in vehicles which were exposed to Nogo-66 longer than or equal to 2 hours (P< 0.05, Fig. 1B). The fluorescence intensity quantitative assay also confirmed the tendency of the p-ILK in N2a cells exposed to Nogo-66 (Fig. 1C, D), however, no significant difference was not observed in the 2h vehicle. For the cells exposed to Nogo-66 (Fig. 1A, B) treatment in variety of time durations, there were no significance differences observed. The results indicated that Nogo-66 suppressed the activation of ILK but had no impact on the expression of total ILK.

**Figure 1.**
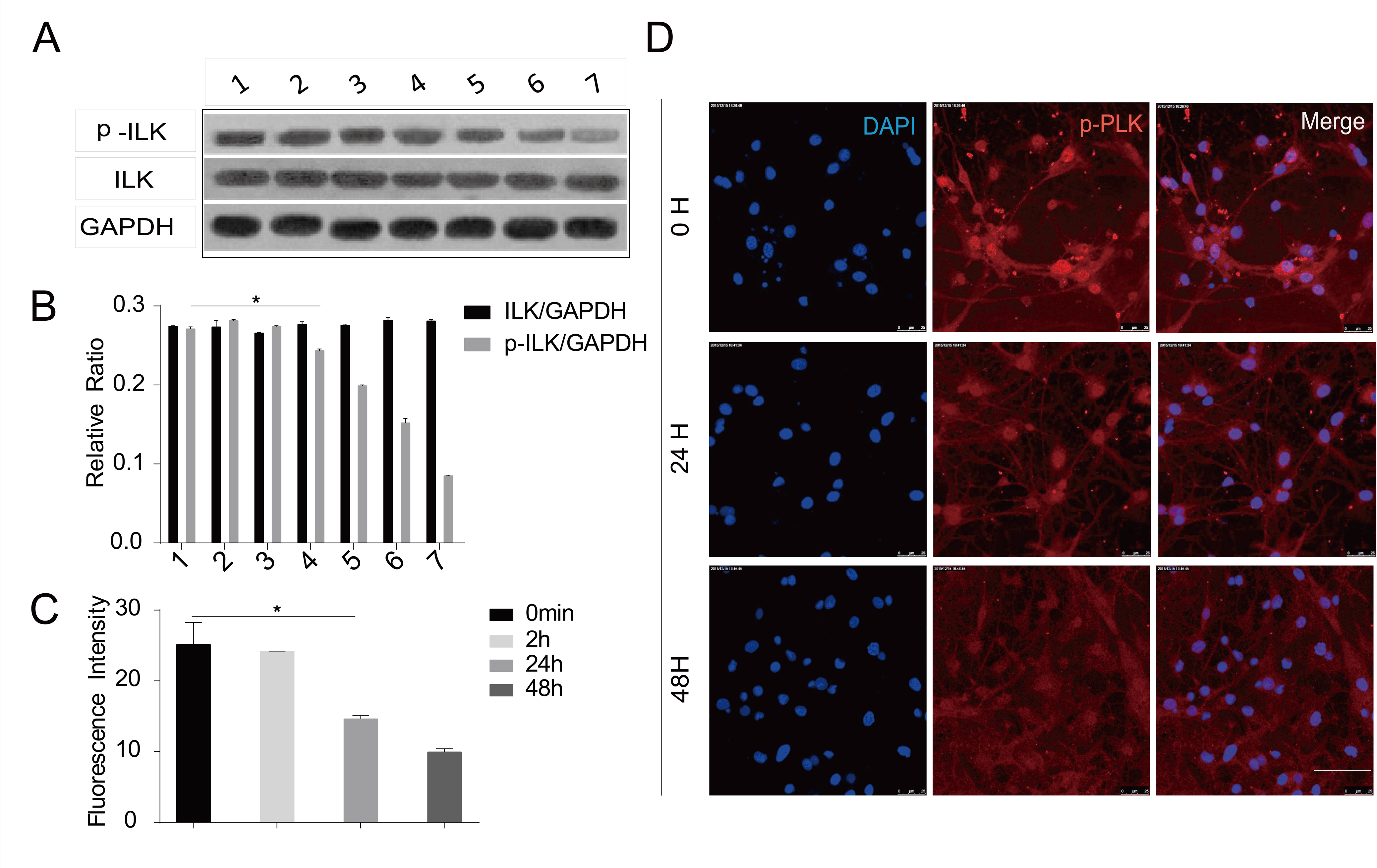
Nogo-66 decreased ILK phosphorylation level in N2a Cells. (A) Treated with Nogo-66 for a staggered time schedule, p-ILK levels in N2a Cells were decreased without apparent change of ILK level. p-ILK at 50kD, ILK at 51kD, GAPDH at 36kD. Quantification of western blotting after Nogo-66 treatment.The experiment was repeated four times. Bands were analyzed by densitometry and values are expressed as a percentage of control (0 min) ± SEM in the three experiments. *P<0.05. (C) Quantification of immunofluorence after Nogo-66 treatment. Immunofluorence densities were analyzed by densitometry and were subtracted by background value for each groupn=4. Error bar indicated mean±SEM. *P < 0.05. (D) Detection of distribution and expression of p-ILK proteins in a high magnification (800×) view in N2a Cells(n=4). Scale bar, 100μm. Values were subtracted by background value for each group.

### Influence of ILK on the expression of tau in N2a Cells

Tau is linked to microtubule dynamics and transport, which play an important role in axonal outgrowth. Previous studies have proven that tau is involved in Nogo-66 mediated signaling pathway of neurite outgrowth downstream inhibition of GSK-3β^[12]^. To investigate the influence of ILK on the expression of tau in N2a Cells treated with Nogo-66 for 24h, manipulation of ILK expression by ILK-Lenti-Ove and shRNA ILK-Lenti-Int treatment was employed(Figure. 2A). The findings demonstrated that the expression of tau was significantly affected by the expression of ILK. Western blot analysis showed that ILK overexpression via ILK-Lenti-Ove significantly elevated the expression level of tau (P < 0.05, comparison between Ove. and control group; Fig. 2D, E), whereas lowering ILK expression with shRNA ILK-Lenti-Int significantly reduced the expression of tau (P < 0.05, comparison between Int. and control group; Fig. 2D, E). Fluorescence intensity quantitative assay was used to confirm the results. (Fig. 2B, C). These findings suggested tau expression was dependent on the level of ILK.

**Figure 2.**
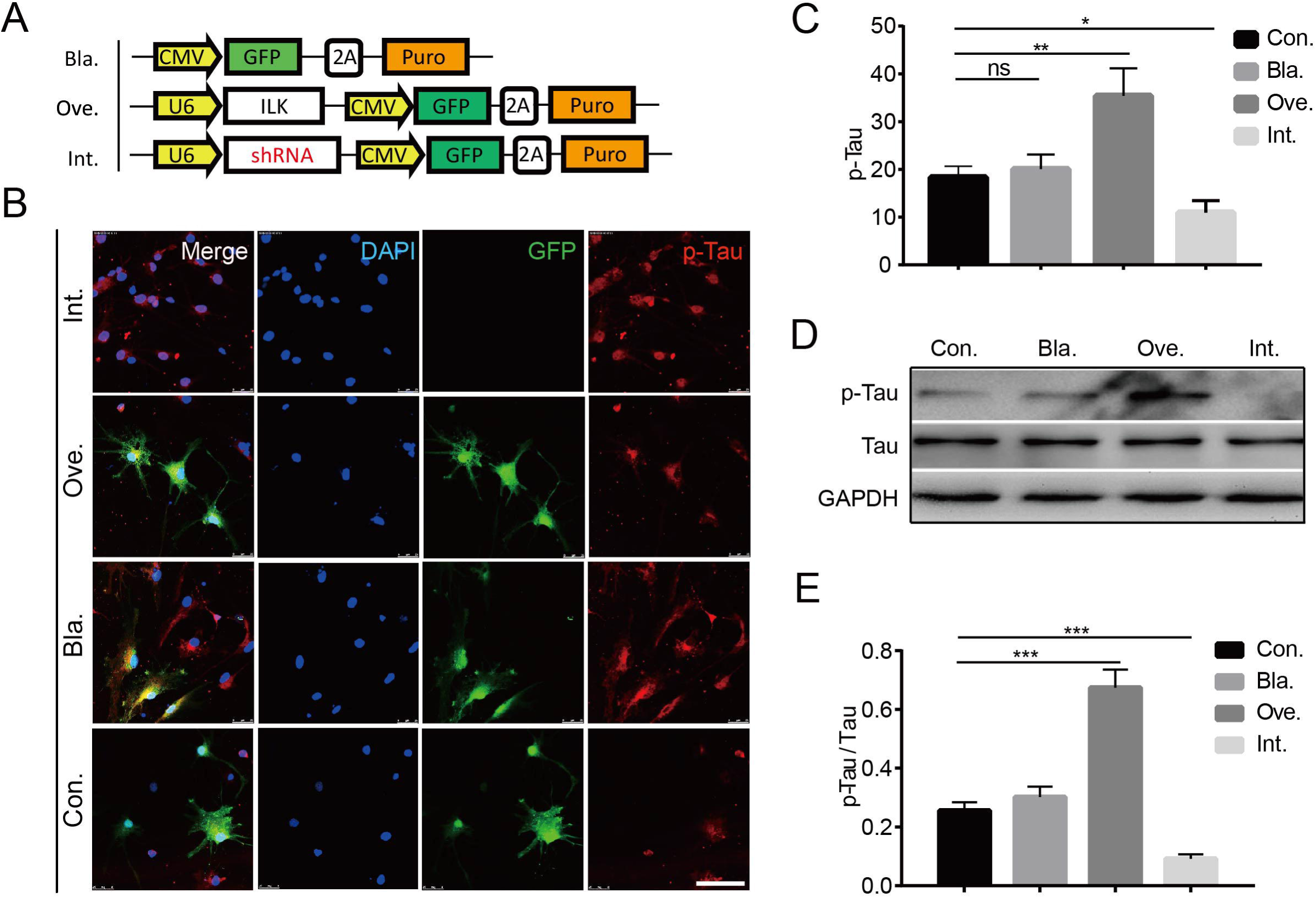
The overexpression of ILK promoted tau phosphorylation level. (A) Schematic illustration of virus construction strategy. Bla, pLenti-EGFP; Ove, pLenti-ILK-EGFP; Int, pLenti-shRNA-EGFP. (B) Tau detected by immunofluorence technicque in N2a Cells in the four groups (n=4). Con, PBS; DAPI, 4‘,6-diamidino-2-phenylindole; GFP, Green Fluorescent Protein. Scale bar, 50um; (C) Quantification of immunofluorence in these four groups after staggered time schedule. Immunofluorence densities were analyzed by densitometry and were subtracted by background value for each group. Error bar indicated mean±SEM. ns represents nonsence; *P < 0.05; **P<0.01. (D) Western-blotting results of tau and p-Tau. (E) Quantification of western blotting(n=4). Bands were analyzed by densitometry and values are expressed as mean±SEM in three experiments. ***P<0.001.

### Influence of ILK on the phosphorylation of PKB and GSK-3β on N2a Cells

Phosphorylation is the way through which kinase acquires its activity, thus p-PKB and p-GSK-3β are the active forms of the kinases. Determining the influence of ILK on the expression levels of p-PKB and p-GSK-3β, this study manipulated ILK expression by ILK-Lenti-Ove and ILK-Lenti-Int treatment. Western blot analysis found that the Ove. Group expressed significantly increased levels of p-PKB at residues Ser473 (Fig. 3A, B) and p-GSK-3β at residues Ser9 (Fig. 4A, B), whereas the Int. group significantly reduced the expression of p-PKB (Fig. 3C, D) and p-GSK-3β (Fig. 4C, D), compared to the control groups. The results illustrated that the activation of ILK was required for the phosphorylation of PKB and GSK-3β.

**Figure 3.**
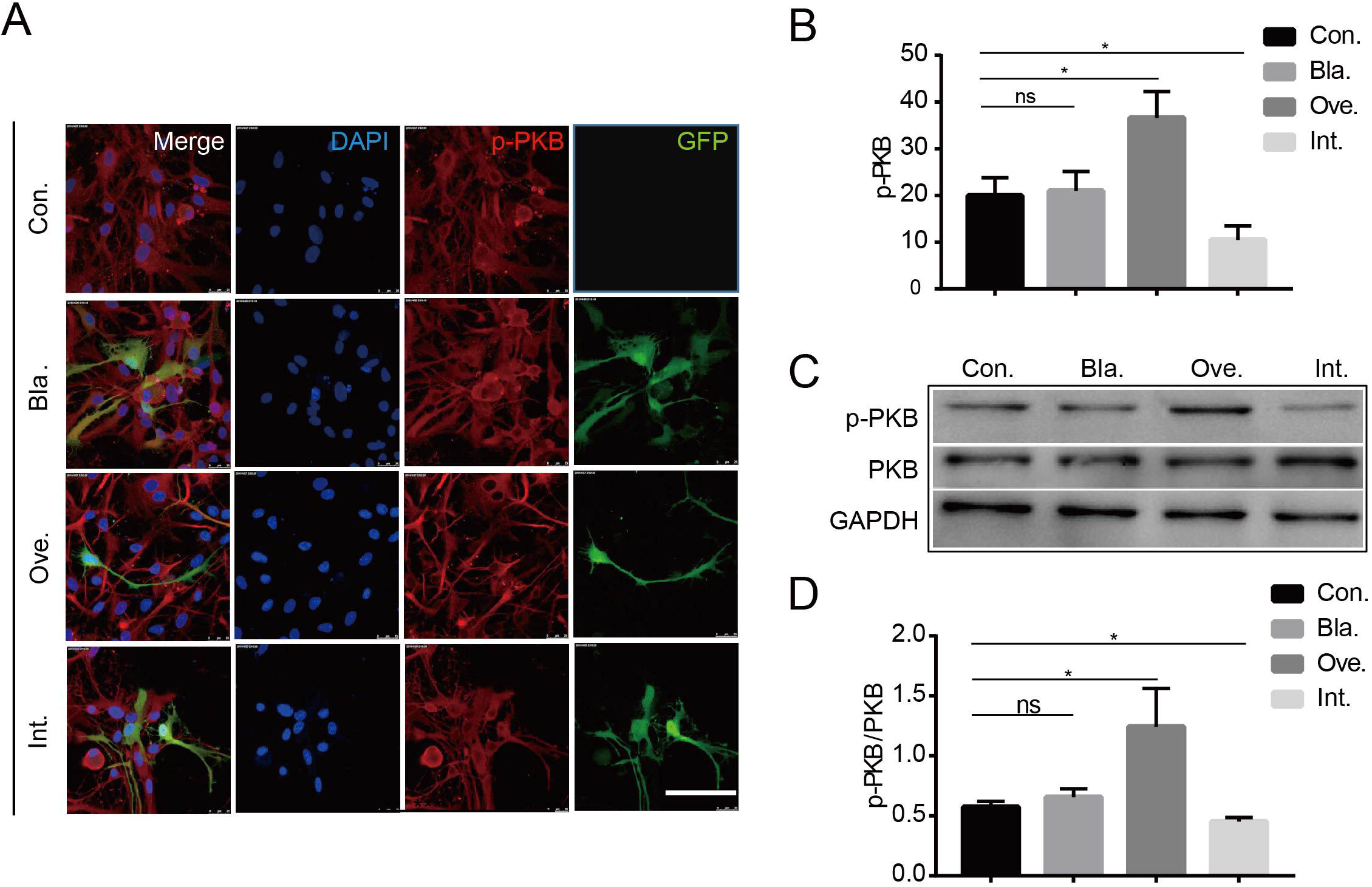
The overexpression of ILK promoted PKB phosphorylation level. (A) p-PKB detected by immunofluorence in N2a cells in four groups. DAPI, 4‘,6-diamidino-2-phenylindole; p-PKB, phosphorylated protein kinase B; GFP, Green Fluorescent Protein. Scale bar, 50um. (B) Quantification of immunofluorence. Immunofluorence densities were analyzed by densitometry and were subtracted by background value for each group. Error bar indicated mean±SEM. *P < 0.05. (C) p-PKB was detected by western-blotting. (D) Quantification of p-PKB via western blotting. Bands were analyzed by densitometry and values are expressed as mean±SEM in three experiments. ns represents nonsense; *P < 0.05.

**Figure 4.**
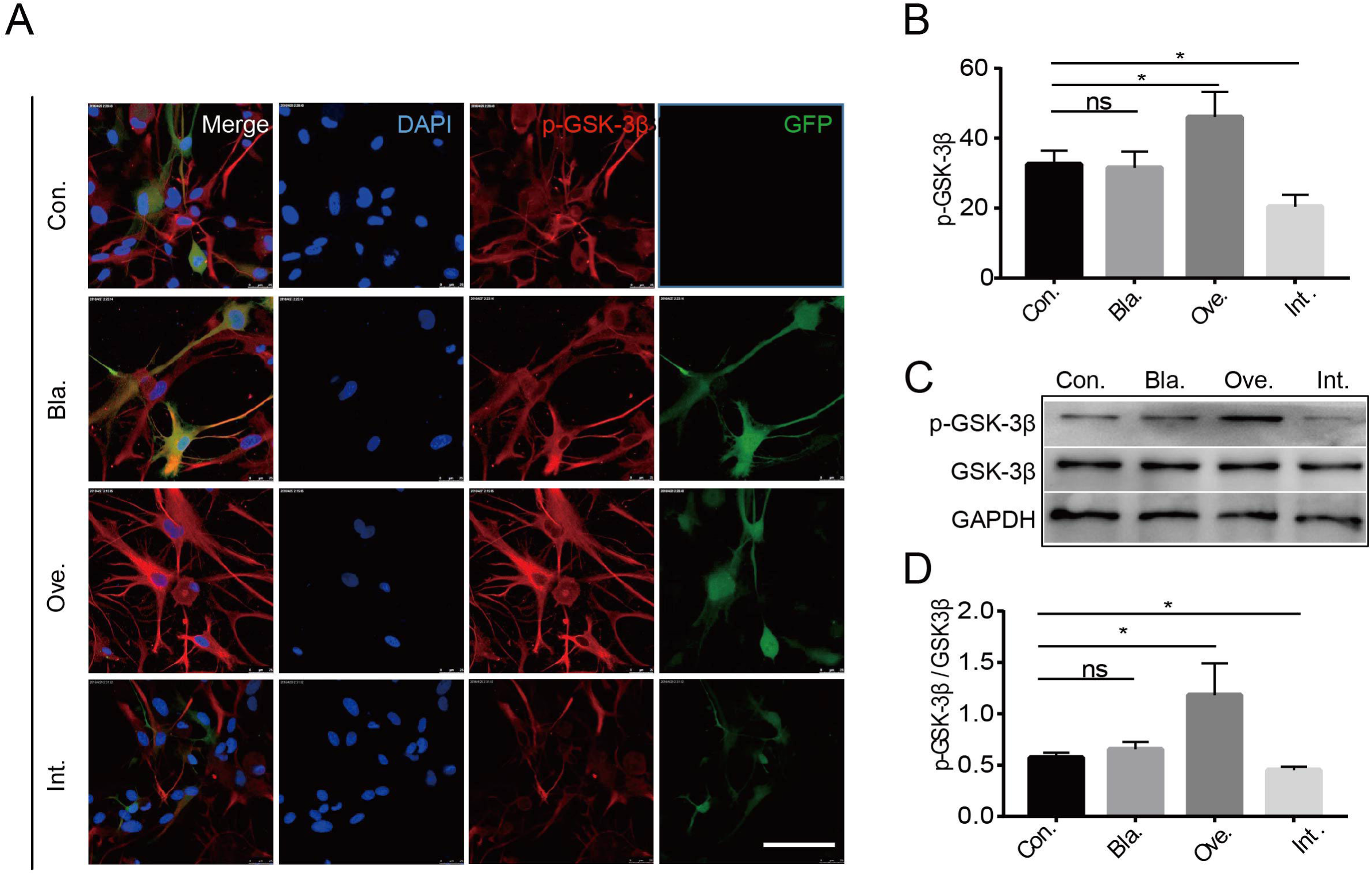
The overexpression of ILK gene promoted GSK-3β phosphorylation level. **(A)** p-GSK-3β detected by immunofluorence in N2a cells. DAPI, 4’,6-diamidino-2-phenylindole; p-GSK-3β, phosphorylated Glycogen Synthase Kinase 3 Beta; GFP, Green Fluorescent Protein. Scale bar, 50um. (B) Quantification of immunofluorence. Densities were analyzed by densitometry and were subtracted by background value for each group(N=10). Error bar indicated mean±SEM. *P < 0.05. p-GSK-3β was detected by western-blotting. (D) Quantification analysis of p-GSK-3β via western blotting(N=4). Bands were analyzed by densitometry and values are expressed as mean±SEM in three experiments. *P < 0.05.

### ILK overexpression ameliorated the inhibitory effect of Nogo-66 in cortical neurons

Neurons were pretreated with EGFP-Lenti (Bla.), ILK-Lenti-Ove and ILK-Lenti-Int, cultivated then in a medium containing Nogo-66 (15 ng/mL) or DMSO for 24h. The neurons were photographed, and the lengths of neurite were measured. Neurite length under normal conditions (Bla. + DMSO) averaged 68.1 ± 12.2μm, while the neurite length shortened to 30.2 ± 7.1 μm in the Bla. + Nogo-66 group, which indicated Nogo-66 significantly inhibited the neurite outgrowth (P < 0.05, Fig. 5). Pretreatment with ILK-Lenti-Ove (Ove. + Nogo-66) significantly ameliorated the inhibitory effect caused by Nogo-66 and kept the neurite length at a similar level (67.5 ± 10.5 μm, P < 0.05 compared to the Bla. + Nogo-66 group). In contrast, pretreatment with ILK-Lenti-Int significantly shortened neurite length to 25.6 ± 6.1 μm (P < 0.05, compared to the Bla. + Nogo-66 group). The results suggested a key role for ILK in the neurite outgrowth inhibition induced by Nogo-66.

**Figure 5.**
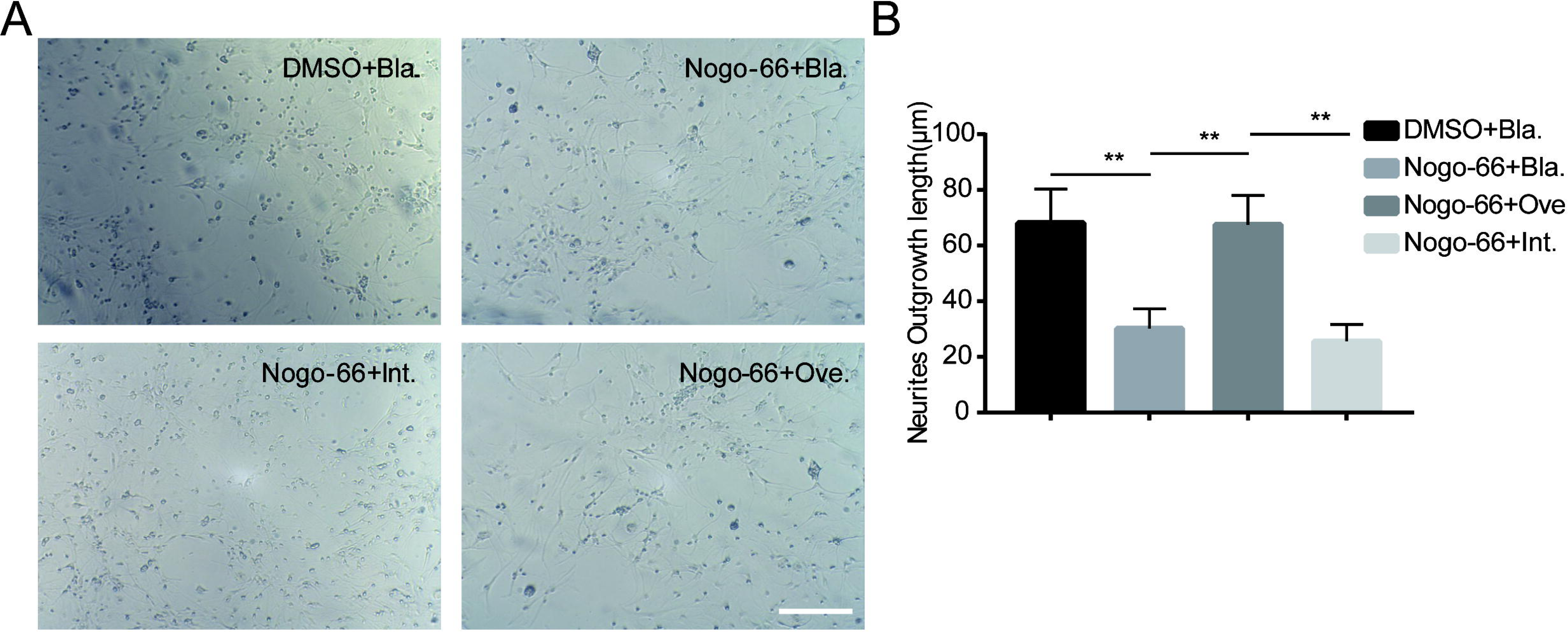
ILK overexpression rescued Nogo-66-induced inhibition of neurite outgrowth in cortical neurons. (A) Neurons images of four groups. (B) The Neurite length of four groupsMeasurements were made on duplicated wells (n= 6). Data were shown as means±SD of three experiments. *P < 0.05.

### ILK overexpression promoted tau expression in complete spinal cord transection model of rats

The expressions of ILK and p-ILK were dynamically observed in the rat model of the complete spinal cord transection. The protein level of ILK held steady after injury, whereas the expression of p-ILK decreased gradually, and the statistical differences were observed after the first day post-operation (Fig. 6A and B). The results were in line with the phenomenon that Nogo-66 inhibited phosphorylation of ILK *in vitro*.

**Figure 6.**
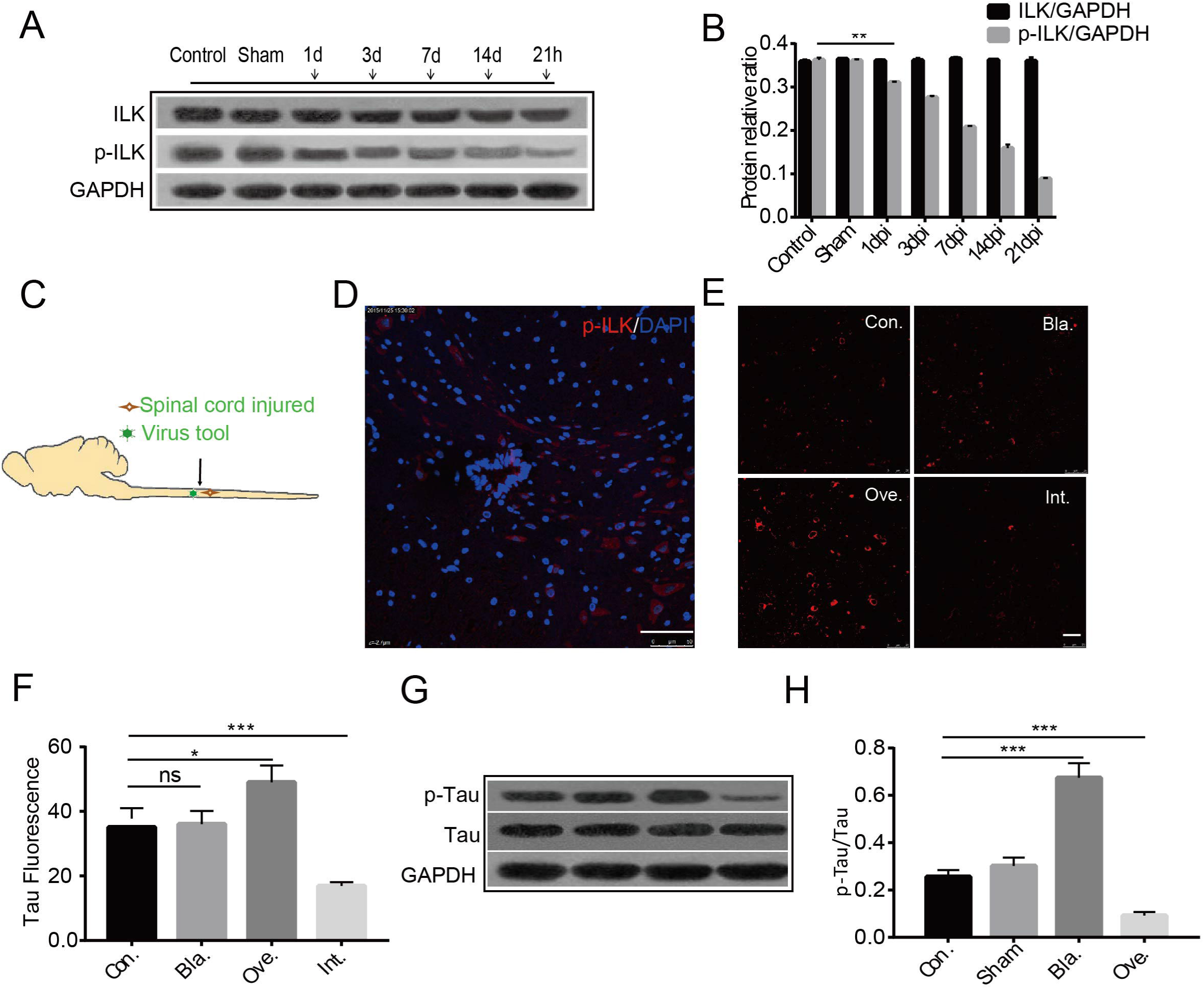
p-ILK level decreased after spinal cord injury and overexpression of ILK in vivo promoted tau phosphorylation level. (A) Western-blotting analysis of the tissue homogenate in spinal cord injured model. The lower line represents GAPDH, the middle line represents p-ILK and the top line represents p-ILK. (B) Quantification of western-blotting analysis in spinal cord injured model. Bands were analyzed by densitometry and values are expressed as mean±SEM in three experiments. *P < 0.05.Cschematic illustration of spinal cord injury and the delivery of lentivirus. Dthe expression of pLenti-ILK-EGFP near injured site of spinal cord. Scale bar, 100um. (E) Protein tau detected by immunofluorence technique in spinal cord injured model with lentivirus injection. Scale bar, 50um. (F) Quantification of immunofluorence in these four groups. *P<0.05.(G) Stereotaxic injection of four groups of vectors into spinal cord and collected the tissue homogenate to make western-blotting analysis. The lower line represents GAPDH, the middle line represents tau, the upper line represents p-tau. (H) Quantification of western-blotting analysis in those four groups. Bands were analyzed by densitometry and values are expressed as mean±SEM in three experiments. *P<0.05.

Determining the influence of ILK on neurite outgrowth *in vivo*, manipulation of ILK expression by ILK-Lenti-Ove and shRNA ILK-Lenti-Int treatment was employed((Fig. 6C and D). The expression of tau, the marker protein of axon, was investigated. The findings demonstrated that the expression of tau (p-S262 tau) was significantly affected by the expression of ILK. Fluorescence intensity analysis showed that overexpression via ILK-Lenti-Ove significantly increased the expression levels of tau (P < 0.05, comparison between Ove. and control group; Fig. 6E, F), whereas lowering ILK expression levels with shRNA ILK-Lenti-Int significantly reduced the expression of tau (P < 0.05, comparison between Int. and control group; Fig. 6E, F). These results were confirmed by Western blotting (Fig. 6G, H). The data indirectly suggested that ILK was involved in neurite outgrowth inhibiting signaling and provided potential targets for the promotion of neurite outgrowth.

## Discussion

Integrins provided a physical link between the cytoskeleton and the extracellular matrix. The clustering of integrins caused the generation of intracellular signals regulating cell growth, survival, differentiation and motility. ILK was discovered as the most prominent interactor interacting with the cytoplasmic domain of β1-integrin subunit^[15]^. Various studies have confirmed that ILK played an important and essential role in regulating cell spreading, cell-cycle progression and the activation of certain number of signaling pathways, through its kinase activity^[16, 17]^.

The current study reports a novel role for the ILK in regulating dendrite growth and yields four major findings. Firstly, the phosphorylation level of ILK can be regulated by Nogo-66. After exposure to the Nogo-66 treatment longer than 2 hours, the activation (phosphorylation) of ILK was significantly suppressed in N2a cells. Secondly, ILK can regulate the phosphorylation of PKB and GSK-3β, and expression level of tau in Nogo-66-treated N2a cells. Research focusing on cancer cells has demonstrated that inhibiting ILK inhibits PKB phosphorylation, and the overexpression or activation of ILK promotes phosphorylation of PKB and cell survival^[18]^. Studies based on neural regeneration found that GSK-3β functioned downstream of ILK to regulate dendrite formation^[19]^. However, to our knowledge, this is the first report on the correlation between ILK and PKB/GSK-3β in Nogo-66 environment. Thirdly, resuscitation of axon elongation was observed in Nogo-66-treated cortical neurons when up-regulating the expression of ILK, indicating that ILK, ameliorated the inhibitory effect of Nogo-66. Finally, after the injection of lentivirus vectors up-regulates the expression of ILK, an increased phosphorylated tau level was observed in the complete spinal cord transection model of rats.

These results imply a crosslink between ILK and Nogo-66 induced neurite outgrowth inhibition signaling pathway. Combining existing evidence and the study’s findings, we put forward the potential mechanism of neurite outgrowth inhibition signaling pathway induced by Nogo-66 as the following: Nogo-66 binds to receptors and then attenuates the phosphorylation and activation of ILK, was presented. Down-activation of ILK leads to decreased levels of p-PKB, which in turn leads to down-expression of p-GSK-3β. Decreased levels of p-GSK-3β danach results in attenuation of phosphorylated tau, which reduces its ability to regulate the stabilization and assembly of microtubules, and eventually causes the inhibition of neurite outgrowth. The presumed mechanism is illustrated in Fig. 7.

**Figure 7.**
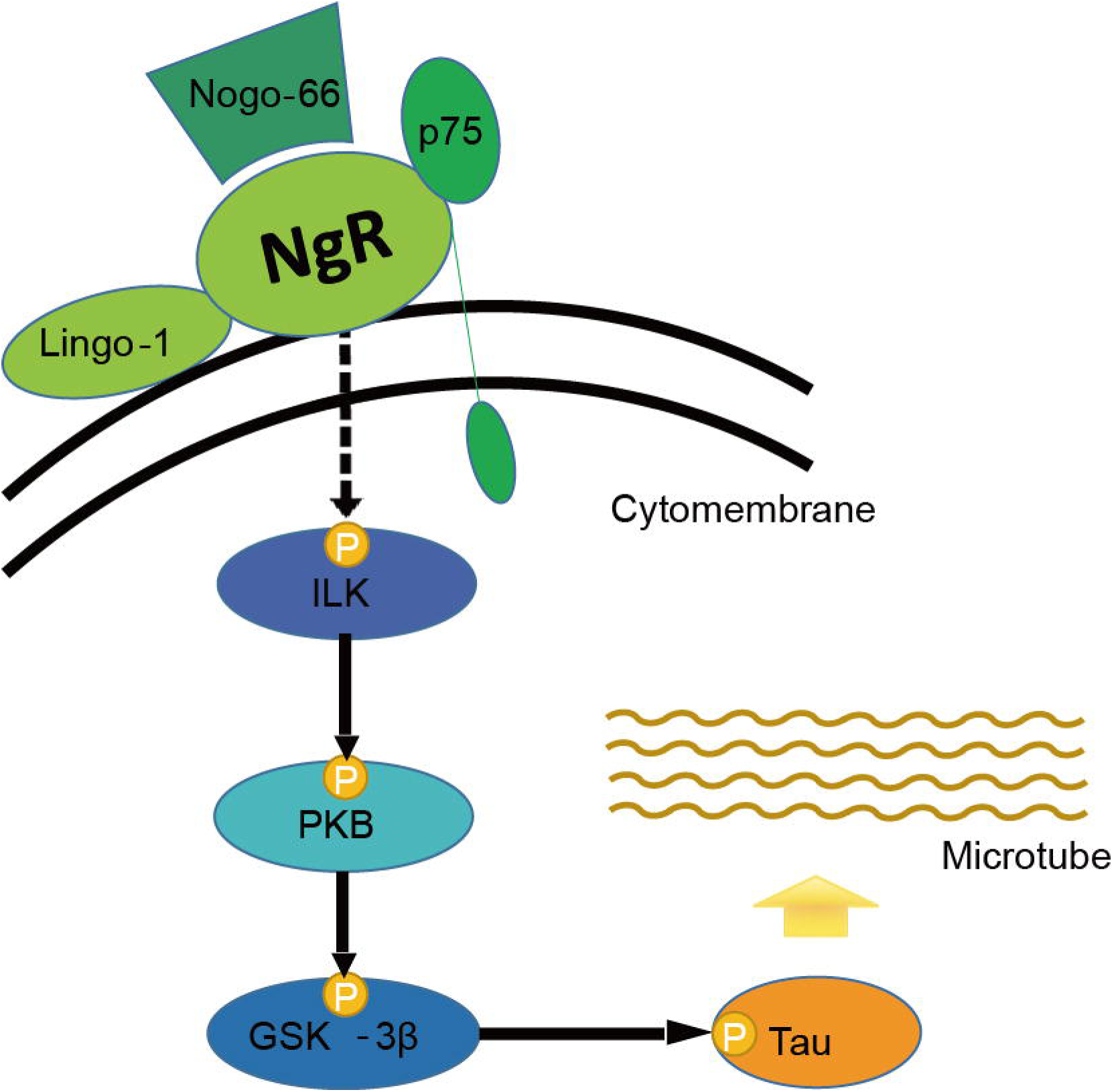
Schematic diagram illustrating presumed mechanism of Nogo-66 induced neurite outgrowth inhibition signaling pathway. Nogo-66 can inhibit ILK phosphorylation. The Level of p-ILK decreases while the total of. ILK actually has no change. Further, downregulated p-PKB caused by decreased p-ILK suppresses the phosphorylation of GSK-3β, with the phosphorylation level of tau further suppressed. Finally the downregulated tau contributing to neurite outgrowth inhibition.

Central nervous system (CNS) is so highly specialized that it cannot effectively rebuild proper circuitry to regenerate after injury. This feature contributes to the failure of the neuron to regenerate and repair after CNS injury^[4, 20]^. Although a range of medication and approaches have been attempted to promote neuron regeneration, the results are still discouraging^[21, 22]^. More and more research continue to validate the critical role that myelin inhibitors (Nogo-A), plays in inhibiting the neurite outgrowth^[23, 24]^. Potential treatments blocking the effects of, such as myelin inhibitors, become the research objectives. This study suggested that the ILK signal is a potential therapeutic target for promoting neurite regeneration.

In this study, successful demonstration that ILK signaling was involved in Nogo-66 induced the inhibition of neurite outgrowth in vitro, were correlated into this study. However, in vivo, primary identification of a crosslink between ILK and tau, a microtubule-associated protein participating in the signaling, without providing direct evidence presenting the relationship between ILK signaling and neurite outgrowth. This was considered to be the flaw of this study. Thus, subsequent research is needed to further confirm the thesis.

In summary, the key role played by ILK in the signaling pathway of Nogo-66-induced inhibition of neurite outgrowth, were reported. The present study further explains the mechanism of Nogo-66 signaling pathway and provides a novel target for promoting neural regeneration.

## Abbreviations

Nogo-A: neurite outgrowth inhibitor
PKB: Protein kinase B
GSK-3β: Glycogen synthase kinase-3β
ILK: Integrin-linked kinase
N2a: Neuro 2a
CNS: central nervous system
OMgp: oligodendrocyte-myelin glycoprotein
MAG: myelin-associated glycoprotein
CSPG: chondroitin sulfate proteoglycan
NgR1: Nogo-66 receptor 1
GADPH: glyceraldehyde-3-phosphate dehydrogenase
SD: Sprague-Dawley
DMEM: Dulbecco’s modified Eagle’s medium
PCR: polymerase chain reaction
SDS-PAGE: sodium dodecyl sulfate-polyacrylamide gel electrophoresis
DMSO: dimetilsulfóxido
PVDF: polyvinylidene difluoride
PBS: polybutylene succinate
SPSS: The Statistical Package for the Social Sciences
SD: standard deviation
ANOVA: One-way analysis of variance

## Disclosure

The authors reported no conflict of interest regarding the materials or methods used in this study or the findings specified in this paper.

## Acknowledgements

This work was supported by the National Natural Science Foundation of China (81671210).

## Author Contributions

Hua Yu, Jian-ying Shen, Peng Fu, Lei Wang, Ye Yuan carried out experiments. Ya-ping Yu and Qiang-ping Wang analyzed the data. Qiang-ping Wang, Nan-xiang Xiong and Hong-yang Zhao designed the experiments. Ya-ping Yu and Qiang-ping Wang wrote the paper. Yi-hao Wang and Hendrik Pool edited the paper.

## Conflict of interest

All authors certify that they have no affiliations with or involvement in any organization or entity with any financial interest in the subject matter or materials discussed in this manuscript.

## Notes

### Competing Interest Statement

The authors have declared no competing interest.

## References

[1] Bradbury EJ, McMahon SB. Spinal cord repair strategies: why do they work? Nat Rev Neurosci 2006, 7: 644–653.

[2] Liu K, Lu Y, Lee JK, Samara R, Willenberg R, Sears-Kraxberger I, et al. PTEN deletion enhances the regenerative ability of adult corticospinal neurons. Nat Neurosci 2010, 13: 1075–1081.

[3] McKerracher L, Winton MJ. Nogo on the go. Neuron 2002, 36: 345–348.

[4] Filbin MT. Myelin-associated inhibitors of axonal regeneration in the adult mammalian CNS. Nat Rev Neurosci 2003, 4: 703–713.

[5] GrandPre T, Li S, Strittmatter SM. Nogo-66 receptor antagonist peptide promotes axonal regeneration. Nature 2002, 417: 547–551.

[6] Viniegra JG, Martinez N, Modirassari P, Hernandez LJ, Parada CC, Sanchez-Arevalo LV, et al. Full activation of PKB/Akt in response to insulin or ionizing radiation is mediated through ATM. J Biol Chem 2005, 280: 4029–4036.

[7] Liu CM, Hur EM, Zhou FQ. Coordinating Gene Expression and Axon Assembly to Control Axon Growth: Potential Role of GSK3 Signaling. Front Mol Neurosci 2012, 5: 3.

[8] Shen JY, Yi XX, Xiong NX, Wang HJ, Duan XW, Zhao HY. GSK-3beta activation mediates Nogo-66-induced inhibition of neurite outgrowth in N2a cells. Neurosci Lett 2011, 505: 165–170.

[9] Wang H, Shen J, Xiong N, Zhao H, Chen Y. Protein kinase B is involved in Nogo-66 inhibiting neurite outgrowth in PC12 cells. Neuroreport 2011, 22: 733–738.

[10] Weingarten MD, Lockwood AH, Hwo SY, Kirschner MW. A protein factor essential for microtubule assembly. Proc Natl Acad Sci U S A 1975, 72: 1858–1862.

[11] Wang Y, Mandelkow E. Tau in physiology and pathology. Nat Rev Neurosci 2016, 17: 5–21.

[12] Zuo YC, Li HL, Xiong NX, Shen JY, Huang YZ, Fu P, et al. Overexpression of Tau Rescues Nogo-66-Induced Neurite Outgrowth Inhibition In Vitro. Neurosci Bull 2016, 32: 577–584.

[13] Mills J, Digicaylioglu M, Legg AT, Young CE, Young SS, Barr AM, et al. Role of integrin-linked kinase in nerve growth factor-stimulated neurite outgrowth. J Neurosci 2003, 23: 1638–1648.

[14] Dedhar S, Williams B, Hannigan G. Integrin-linked kinase (ILK): a regulator of integrin and growth-factor signalling. Trends Cell Biol 1999, 9: 319–323.

[15] Hannigan GE, Leung-Hagesteijn C, Fitz-Gibbon L, Coppolino MG, Radeva G, Filmus J, et al. Regulation of cell adhesion and anchorage-dependent growth by a new beta 1-integrin-linked protein kinase. Nature 1996, 379: 91–96.

[16] Yamaji S, Suzuki A, Kanamori H, Mishima W, Yoshimi R, Takasaki H, et al. Affixin interacts with alpha-actinin and mediates integrin signaling for reorganization of F-actin induced by initial cell-substrate interaction. J Cell Biol 2004, 165: 539–551.

[17] Persad S, Dedhar S. The role of integrin-linked kinase (ILK) in cancer progression. Cancer Metastasis Rev 2003, 22: 375–384.

[18] Persad S, Attwell S, Gray V, Delcommenne M, Troussard A, Sanghera J, et al. Inhibition of integrin-linked kinase (ILK) suppresses activation of protein kinase B/Akt and induces cell cycle arrest and apoptosis of PTEN-mutant prostate cancer cells. Proc Natl Acad Sci U S A 2000, 97: 3207–3212.

[19] Naska S, Park KJ, Hannigan GE, Dedhar S, Miller FD, Kaplan DR. An essential role for the integrin-linked kinase-glycogen synthase kinase-3 beta pathway during dendrite initiation and growth. J Neurosci 2006, 26: 13344–13356.

[20] York EM, Petit A, Roskams AJ. Epigenetics of neural repair following spinal cord injury. Neurotherapeutics 2013, 10: 757–770.

[21] Finelli MJ, Wong JK, Zou H. Epigenetic regulation of sensory axon regeneration after spinal cord injury. J Neurosci 2013, 33: 19664–19676.

[22] Zhang C, Tu F, Zhang JY, Shen L. E-cadherin-transfected neural stem cells transplantation for spinal cord injury in rats. J Huazhong Univ Sci Technolog Med Sci 2014, 34: 554–558.

[23] Kottis V, Thibault P, Mikol D, Xiao ZC, Zhang R, Dergham P, et al. Oligodendrocyte-myelin glycoprotein (OMgp) is an inhibitor of neurite outgrowth. J Neurochem 2002, 82: 1566–1569.

[24] Schwab ME. Functions of Nogo proteins and their receptors in the nervous system. Nat Rev Neurosci 2010, 11: 799–811.

